# Fast weight recovery, metabolic rate adjustment and gene-expression regulation define responses of cold-stressed honey bee brood

**DOI:** 10.1101/2020.06.15.152389

**Authors:** Leonor Ramirez, Facundo Luna, Claudio Andoni Mucci, Lorenzo Lamattina

**Affiliations:** Instituto de Investigaciones Biológicas (IIB), Consejo Nacional de Investigaciones Científicas y Técnicas (CONICET) – Universidad Nacional de Mar del Plata (UNMdP), CC1245, 7600 Mar del Plata, Argentina; Laboratorio de Ecología Fisiológica y del Comportamiento, Instituto de Investigaciones Marinas y Costeras (IIMyC), CONICET – UNMdP, CC1245, 7600 Mar del Plata, Argentina

## Abstract

In temperate climates, low ambient temperatures in late winter and in spring can result in cold stress conditions in brood areas of weakened honey bee colonies, leading to increased levels of developmental interruptions and death of the brood. Very little is known about the physiological and molecular mechanisms that regulate honey bee brood responses to acute cold-stress. Here, we hypothesized that central regulatory pathways mediated by insulin/insulin-like peptide signalling (IIS) and adipokinetic hormone (AKH) are linked to metabolic changes in cold-stressed honey bee brood. *A. mellifera* brood reared at suboptimal temperatures showed diminished growth rate and arrested development progress. Notably, cold-stressed brood rapidly recovers the growth in the first 24 h after returning at control rearing temperature, sustained by the induction of compensatory mechanisms. We determined fast changes in the expression of components of IIS and AKH pathways in cold-stressed brood supporting their participation in metabolic events, growth and stress responses. We also showed that metabolic rate keeps high in brood exposed to stress suggesting a role in energy supply for growth and cell repair. Additionally, transcript levels of the uncoupling protein MUP2 were elevated in cold-stressed brood, suggesting a role for heat generation through mitochondrial decoupling mechanisms and/or ROS attenuation. Physiological, metabolic and molecular mechanisms that shape the responses to cold-stress in honey bee brood are addressed and discussed.

## INTRODUCTION

Temperature is an environmental factor that has a dramatic effect on survival, growth and development in ectotherms like honey bee larvae, if the heating capacity of the colony is compromised (1–3). Precise control of nest temperature between 33°C and 36°C can be regarded as one of the major innovations in honeybee biology made possible by the evolution of their societies (4). The main active process achieved for thermal homeostasis of honey bee colonies is the so-called ‘‘endothermy on demand’’ of adults (5). Under cold-stress perception, adult bees increase the heat production with the thoracic flight muscles, resulting in a high number of endothermic individuals, especially in the brood nest (5). Other measures like migration activity within the colony, evaporative cooling, and regulation of internal heat transport via convection (fanning) are used by honey bees (6).

Honey bee brood rearing starts in winter and peaks in spring, decreases through summer, and ceases in early fall in temperate climates (7). Cold snaps or periods of extreme temperatures for three or more consecutive days frequently occur during winter and early spring in temperate climates. Therefore, under these conditions, a deficient colony thermoregulation is a significant source of cold stress for honey bee brood. This is a subject to take into account due to climate change is predicted to cause increasing temperature variability, including cold snaps (8). Even if workers heat the brood to keep it at a constant temperature, other stressors contribute to affect the heating of the nest, brood and the colony health (9). Different stressors are contributing to honey bee colony failure, most with a strong anthropogenic component (10). Managed honey bee stocks were reported to decline in North America, many European countries and in South America (10,11). Perry et al. (12) showed that when forager death rates increase, an increasingly younger forager force causes an accelerated population decline in the colony. This results in a decline of the adult bee population and loss of colony thermal homeostasis with a consequent brood exposure to suboptimal temperatures.

While there is a detailed knowledge of the colony’s strategies to control brood nest temperature, less is known about how honey bee brood responds when grow at low temperatures. Recently, we showed that survival diminishes approximately 60% in *in vitro*-reared *Apis mellifera* brood kept at 25°C for 3 days, compared to brood reared at the standard temperature 34°C (13). Moreover, we demonstrated that, while honey bee brood reared at 34°C reaches adult emergence in approximately 21 days after larvae hatching, there was a significant delay in development of cold-stressed brood, where adults emerged at day 25 (13). The timing of brood onset is critical for colony fitness as it influences the ability to exploit spring bloom (14). Stressful conditions during developmental processes cause not only developmental arrest, but also consequences on bee adults (1–3). The physiological mechanisms behind the cold sensitivity of honey bee brood and the compensatory counteractions should be studied in more detail in order to understand their phenotypic plasticity. This knowledge will provide insights to act on and to prevent undesirable effects on adult honey bee life and colony health.

Here, we predict that central regulatory pathways, involved in anabolic and catabolic processes, are linked to metabolic alterations in cold-stressed honey bee brood and counteract the deleterious effects of stress. We focused on the insulin/insulin-like peptide signalling (IIS) which controls growth, nutrient administration and body size (15), the neuropeptide Adipokinetic hormone (AKH) that mobilizes lipids and carbohydrates in stressed conditions (16), and also the mitochondrial responses through the uncoupling protein (UCP) (17). Changes in body mass, metabolic rate, ATP levels and carbohydrate content were correlated with gene expression profile from cold-stressed honey bee brood. Our results provide new insights on the physiological adjustments driven by gene expression and metabolic responses to cope with cold-stress in honey bee brood. Moreover, we analysed the critical recovery process after the stress based on compensatory resources required to reinitiate the growth processes.

## MATERIAL AND METHODS

### Rearing of larvae

Larvae of *Apis mellifera* (*A. mellifera ligustica* - *A. mellifera mellifera*) of some hours from hatching were collected from combs of colonies of the experimental apiary belonging to the Centro de Investigación en Abejas Sociales (CIAS) located in Santa Paula, route 226, Km 10, Mar del Plata, Argentina. Larvae were transferred to rearing plates and were incubated under standardized conditions at the laboratory (13,18,19). The rearing conditions were maintained at 34°C and 96% humidity during the larvae stage and 34°C and 70% humidity during the prepupae/pupa stage. The amount and composition of the diet of honey bee larvae were as detailed in Ramirez et al. (13) (S1 Table). A group of larvae reared 4 days at 34°C was exposed to cold stress by incubating it at 25°C as described in a previous work (13) (and then returned to 34°C until they stopped growth, see S1 Fig). In our experiment, honey bee brood was not assisted by nurse bees and could not escape from the 25°C cold stress as we previously described (13). Body mass was measured in individuals from 4-days-old until 12-days-old, after dabbing them off with filter paper. Brood pictures were taken with a digital camera attached to a stereomicroscope. Magnification (sufficient to nearly fill the microscope field of view) was held constant for each structure.

### Metabolic Rate

Respirometric technique (20) was used to measure metabolic rate (MR). Carbon dioxide production was measured using a computerized positive pressure open-flow respirometry system (Sable Systems, Las Vegas, USA). Honey bee brood was individually placed in a small cylindrical chamber (30 mL) that received air at 80 mL min^−1^ from a flowmeter (Sierra Systems; standard temperature and pressure corrected). Air passed through a CO_2_-absorbent (Sodalime, Sigma-Aldrich) and a water scrubber (Drierite, W.A. Hammond Drierite Co.) before going through the chamber.

The temperature was established by placing the chamber in a temperature-regulated incubator (±0.1°C, SIMEDIX). Excurrent air from the chamber was moved through a CO_2_ analyzer (CA-10; Sable Systems, Las Vegas, USA). CO_2_ analyzer was calibrated using standard procedures. First, zero-point calibration was performed using CO_2_ free gas. Second, spam calibration was implemented using a spam gas. Each honey bee larva was measured at 34 o 25°C according the rearing condition during 30-40 min but only the last 15-20 min period were considered for MR estimation. Data were captured and processed using a UI2 interface and Expedata software (Sable Systems) and converted into μL CO_2_ min^−1^ using Expedata. Body mass (mg) was measured before each experiment.

### Determination of ATP

Each individual was washed with 1X phosphate buffered saline (PBS; 0.43 mM Na_2_HPO_4_, 0.14 mM KH_2_PO_4_, 13.7 mM NaCl, 0.27 mM KCl), dried on filter paper to remove food debris and homogenized with 1X reaction buffer (500 mM Tricine buffer, pH 7.8, 100 mM MgSO_4_, 2 mM EDTA and 2 mM NaN_3_). The supernatant was harvested by centrifugation at 10000 xg for 15 min at 4°C. ATP content was determined with an ATP determination kit (Molecular Probes, Invitrogen, Thermo Fisher Scientific), following the manufacturer’s instruction. Supernatant aliquots (10 μL each of a 1:100 dilution) were added to 90 μL of reaction buffer in each well of a 96-well plate. Luminescence was measured using a Luminoskan Ascent (Thermo Fisher Scientific). ATP content in the experimental samples was calculated from the ATP standard curve.

### Quantification of transcript levels by real time PCR (qPCR)

Total RNA from individuals was extracted with TRIzol reagent (Invitrogen, Thermo Fisher Scientific) as described by the manufacturer’s instruction. One microgram of RNA was reverse transcribed into cDNA using oligo-dT and MMLV-RT (Moloney Murine Leukemia Virus Reverse Transcriptase, Invitrogen, Thermo Fisher Scientific). Before cDNA synthesis, genomic DNA in the samples was digested with DNAsa (Ambion, Thermo Fisher Scientific) according to the manufacturer’s instructions. The cDNA was used as template for real-time PCR analysis. Reactions were performed on a Step-one Real-time PCR machine from Applied Biosystems (California, USA) with Fast Universal SYBR Green Master Rox (Roche, Merck). The primer sequences are listed in S2 Table. To verify the purity of the PCR products, a melting curve was produced after each run. LinRegPCR was the program employed for the analysis of real time RT-PCR data (21). The transcript relative quantification results were determined from the ratio between the starting concentration value of the analysed mRNAs and the reference Actin mRNA in each sample. The mean and standard error was calculated from values of the transcript quantification obtained in each biological replicate.

### Statistics

All data are presented as mean ± SEM, except for S2 Fig where data is presented in box-plots. Gene expression and ATP content data were log transformed to approximate assumption of normality as verified by Shapiro and Levene’s homogeneity test. All the others datasets of our study achieved assumptions of normality without transformation. ANOVA was used to test for differences between body mass, metabolic rate, and specific molecular variables, such as ATP content and gene expression, and post-hoc was performed by Tukey tests. Data were analysed by linear mixed models with the R package “nlme” (22) in R (23) inside RStudio (24). The “rearing temperature” (34°C and 25°C) and “brood age” (days after larva hatching) were included in the model as fixed effects and “the independent experiment” was included as random effects. All statistical tests used α=0.05 to establish significance.

## RESULTS

The growth curve of honey bee brood exposed to cold-stress (25°C) for 3 days strongly differed compared to brood reared continuously at control temperature (34°C) (Fig 1A). During the cold-stress period at 25°C the body mass increase of honey bee brood was less than 2 times compared to brood continuously incubated at 34°C (*t*_566_=23.3, *P*<0.0001; S2A Fig). Honey bee brood grown at 34°C gained body mass until day 6 after larva hatching while in cold-stressed brood the gaining weight and larval period was extended until day 9 of development, 2 days after they had been transferred back to control temperature (Fig 1A and B). The maximum body mass observed in 9-days-old cold-stressed brood was similar to that observed in 6-days-old brood continuously incubated at 34°C (ANOVA, *F*_1_,_127_=1.3, *P*=0.2564; S2B Fig), showing that compensatory growth occurred in cold-stressed brood transferred back to control temperature.

**Fig 1.**
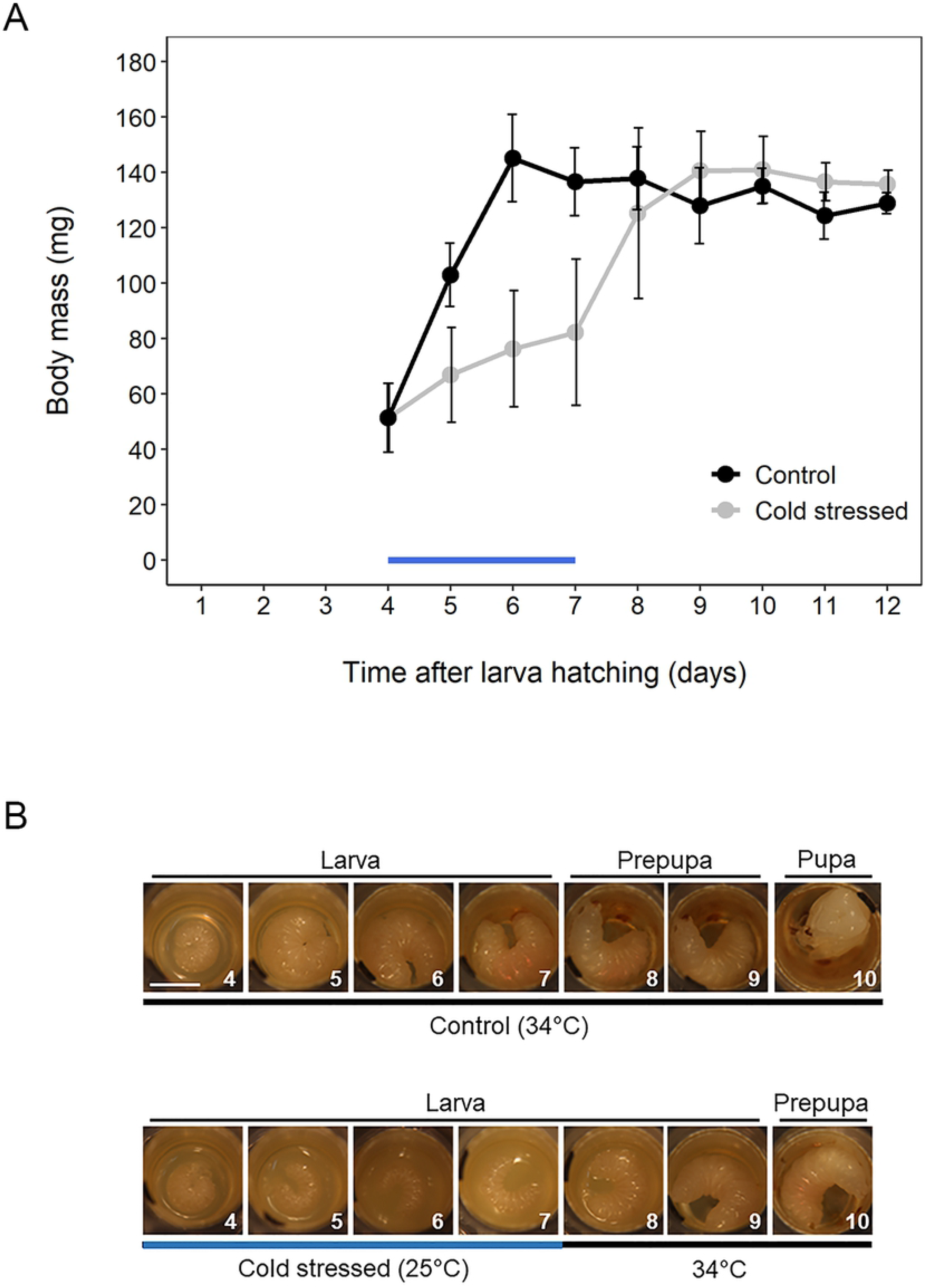
Body mass of cold-stressed honey bee brood through growth in comparison to untreated controls. Four-day-old larvae were reared continuously at control temperature (34°C) or under cold stress (25°C) for 3 days and then were returned to control conditions. A: Growth of individuals incubated at 34°C (black circles) and individuals exposed to cold stress (grey circles). B: Representative pictures of honey bee brood reared at control or cold stressed conditions where the different developmental stages are indicated. Blue line indicates the duration of cold stress.

In cold-stressed brood, the food accumulated in rearing wells, indicating that food intake decreased in these larvae (S3 Fig). This fact correlates with the diminished growth rate of brood exposed to cold-stress.

To understand the physiological processes associated with cold-stress response in honey bee brood, we measured the metabolic rate (MR). Changes occured in the MR mediated by the developmental stage and the current rearing temperature (ANOVA, interaction effect between rearing temperature and brood age, *F*_5,160_=68.6, *P*<0.0001; Fig 2). When honey bee brood was continuously reared at 34°C, the MR gradually decreased from day 4 to day 9 after larva hatching (*t*_11_=8.5, *P*=0.0001; Fig 2). The MR in the cold-stressed larvae decreased during the first 48 h of low temperature (4-days-old vs 6-days-old: *t*_161_=13.9, *P*<0.0001; Fig 2), at the same amount as the control larvae reared at 34°C (6-days-old cold-stressed vs control brood: *t*_161_=−1.7, *P*=0.8689; Fig 2). Then, while MR remained at very low levels in control brood it increased in cold-stressed brood on day 8 (8-days-old cold-stressed vs control brood: *t*_161_=19.2, P<0.0001; Fig 2).

**Fig 2.**
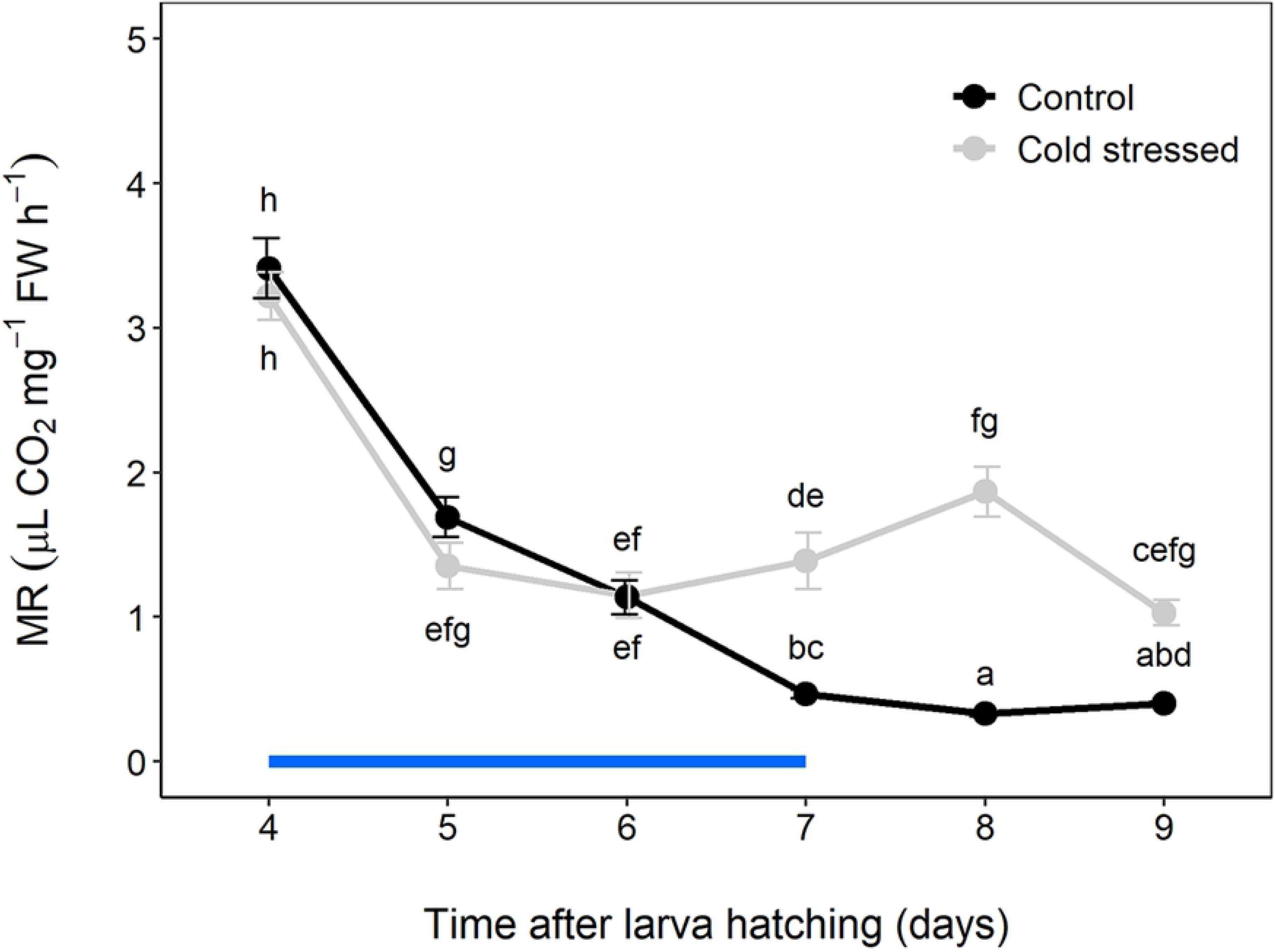
Metabolic rate of cold-stressed honey bee brood in comparison to untreated controls. Four-day-old larvae were reared continuously at control conditions (34°C) or under cold stress (25°C) for 3 days and then were returned to control conditions. Metabolic rate was measured employing a respirometer as detailed in Material and Methods. Data is shown as mean ± SD. Significant differences between honey bee brood incubated at control (black line) or cold stressed (grey line) are indicated with different letters (ANOVA followed by post-hoc comparisons with Tukey). Blue line indicates the duration of cold stress.

To get insights into changes in cellular energy levels under cold stress, the ATP amount was determined in honey bee brood. ATP levels significantly increased 24 h after initiating the cold stress in honey bee brood (5-days-old cold-stressed vs control brood: *t*_105_=6.1, *P*<0.0001; S4 Fig). ATP levels in brood continuously reared at 34°C increased on day 7 after hatching (4-days-old vs 7-days-old control brood: *t*_105_=−4.4, *P*=0.0008; S4 Fig). When cold-stressed brood was returned to 34°C, ATP levels trended to diminish after 24 h, but this was not statically significant (7-days-old vs 8-days-old cold-stressed brood: *t*_105_=2.7, *P*=0.1314; S4 Fig).

Uncoupling proteins (UCPs) mediate mitochondrial thermogenesis in mammals. In *Drosophila melanogaster*, DmUCP4C has been suggested to be an UCP associated with energy dissipation, being essential for larval development at low temperatures (25). In *A. mellifera*, we found two genes with high similarity to the *DmUCP4C* gene, *MUP1* and *MUP2*. Transcript levels of *MUP1* were higher in 8 and 9-days-old cold-stressed brood than in brood continuously reared at 34°C (8-days-old: *t*_141_=4.7, *P*=0.0005; 9-days-old: *t*_141_=4.3, *P*=0.0028; Fig 3A). On the other hand, transcript levels of *MUP2* decreased from day 4 to day 7 in brood reared at 34°C (*t*_175_=9.6, *P*<0.0001; Fig 3B), whereas their levels were kept elevated in cold-stressed brood (*t*_175_=2.3, *P*=0.56; Fig 3B), even one day after returning to 34°C (*t*_175_=3.1, *P*=0.1229; Fig 3B). *MUP2* expression followed a similar pattern to that determined for the MR, suggesting a relationship between these two parameters.

**Fig 3.**
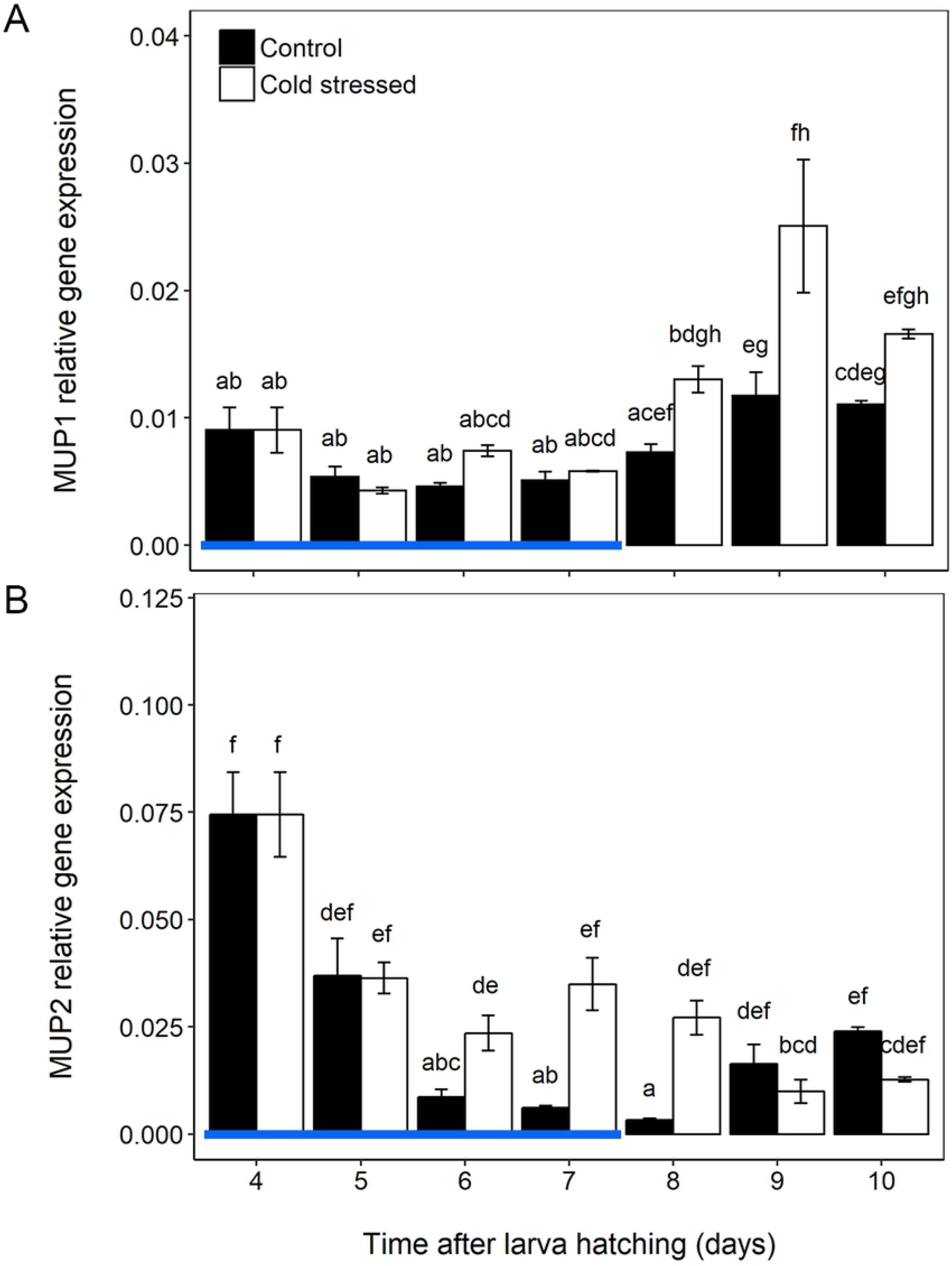
Transcript levels of candidate genes of mitochondrial uncoupling proteins (MUP) in response to cold stress in honey bee brood. Four-day-old larvae were reared continuously at control conditions (34°C) or under cold stress (25°C) for 3 days and then were returned to control conditions. The relative gene expression of MUP1 (A) and MUP2 (B) were determined by RT-qPCR using Actin as a reference gene for normalization. The data are shown as the mean ± SD. Significant differences between control (black boxes) and stressed (white boxes) honey bee brood are indicated with different letters (ANOVA followed by post-hoc comparisons with Tukey). Blue line indicates the duration of cold stress.

The insulin/insulin-like peptide signalling (IIS) regulates physiological processes in insects, particularly growth and stress responses (26). In *D. melanogaster* the IIS pathway is activated in larvae reaching a critical weight leading to the inhibition of the transcription factor Forckhead Box class O (FoxO) that arrests the development and avoids proceeding to the metamorphosis (27). Fig 4A shows a simplified scheme of that pathway. Growth and development are severely affected in cold-stressed brood, thus we analysed if the transcript levels of IIS components Insulin like Peptide 1 (ILP1), Insulin like Peptide 2 (ILP2), Insulin Receptor 1 (InR1) and FoxO are regulated by low temperature. Transcript levels of *ILP1* were not affected in cold-stressed larvae compared to brood continuously reared at 34°C (ANOVA, *F*_1,228_=1.2, *P*=0.2682; Fig 4B). In contrast, while transcript levels of *ILP2* increased constantly from day 4 to day 10 after larva hatching in brood continuously reared at 34°C (*t*_229_=−11.5, *P*<0.0001; Fig 4C), it remained at very low levels in cold-stressed brood (5-days-old: *t*_229_=−3.5, *P*=0.0359; 6-days-old: *t*_229_=−7.1, *P*<0.0001; 7-days-old: *t*_229_=−10.5, *P*<0.0001; 8-days-old: *t*_229_=−8.7, *P*<0.0001; 10-days-old: *t*_229_=−8.4, *P*<0.0001; Fig 4C). Two days after returning to control temperature, the 9-days-old cold-stressed brood reached the same levels of *ILP2* transcript than non-stressed brood (*t*_215_=−0.6, *P*=0.99; Fig 4C). Coincidently, at that time, cold-stressed brood reached the maximum body mass (Fig 1A).

**Fig 4.**
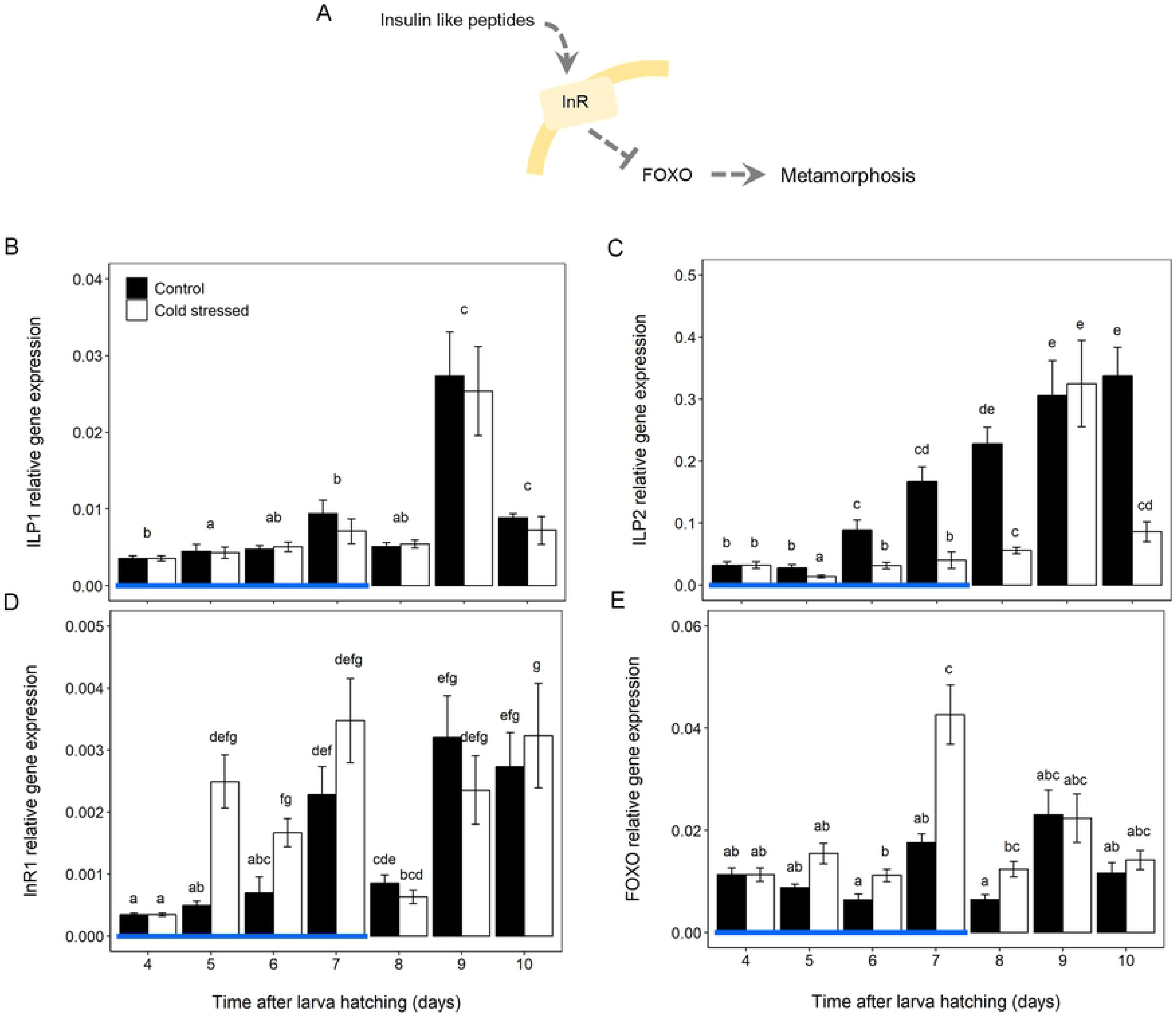
Transcript levels of IIS pathway components in response to cold stress in honey bee brood. Four-day-old larvae were reared continuously at control conditions (34°C) or under cold stress (25°C) for 3 days and then were returned to control conditions. A: Simplified model showing the signalling cascade involving different components of IIS pathway. B-E: Relative gene expression of insulin like peptide 1 (ILP1, B), insulin like peptide 2 (ILP2, C), Insulin receptor 1 (InR1, D) and Forkhead Box class O (FOXO, E) were determined by RT-qPCR using Actin as a reference gene for normalization. The data are shown as the mean ± SD. Significant differences between control (black bars) and stressed (white bars) honey bee brood are indicated with different letters (ANOVA followed by post-hoc comparisons with Tukey). Blue line indicates the duration of cold stress.

Regarding the expression of *InR1*, a slight increase was observed during the cold-stress period compared with brood reared at control temperature (5-days-old: *t*_210_=7.6, *P*<0.0001; 6-days-old: *t*_210_=7.5, *P*<0.0001; Fig 4D). The transcript levels of *FoxO* were not affected in cold-stressed brood compared to those reared at control temperatures at early or late times after larva hatching (5-days-old: *t*_205_=3.3, *P*=0.069; 9-days-old: *t*_205_=−0.2, *P*=0.99; 10-days-old: *t*_205_=2.1, *P*=0.6733; Fig 4E). However, an increase was detected in 6, 7 and 8-days-old cold-stressed brood (6-days-old: *t*_205_=4.8, *P*=0.0003; 7-days-old: *t*_205_=4.6, *P*=0.0006; 8-days-old: *t*_205_=5.3, *P*<0.0001; Fig 4E).

We also analysed the transcript levels of *AKH* and its receptor *AKHR*. Almost no differences were observed for the AKH hormone transcript between control and cold-stressed brood (10-days-old: *t*_224_=3.9, *P*=0.0072; Fig 5A). On the contrary, while *AKHR* expression levels steadily decreased from day 4 to day 8 in brood reared at 34°C (4-days-old vs 8-days-old: *t*_186_=7.3, *P*<0.0001; Fig 5B), its levels remained high in cold-stressed brood (4-days-old vs 8-days-old: *t*_186_=−0.9, *P*=0.9993; Fig 5B). *AKHR* transcript levels abruptly diminished two days after brood had been transferred from cold-stress to control temperature (8-days-old vs 9-days-old: *t*_186_=6.6, *P*<0.0001; Fig 5B).

**Fig 5.**
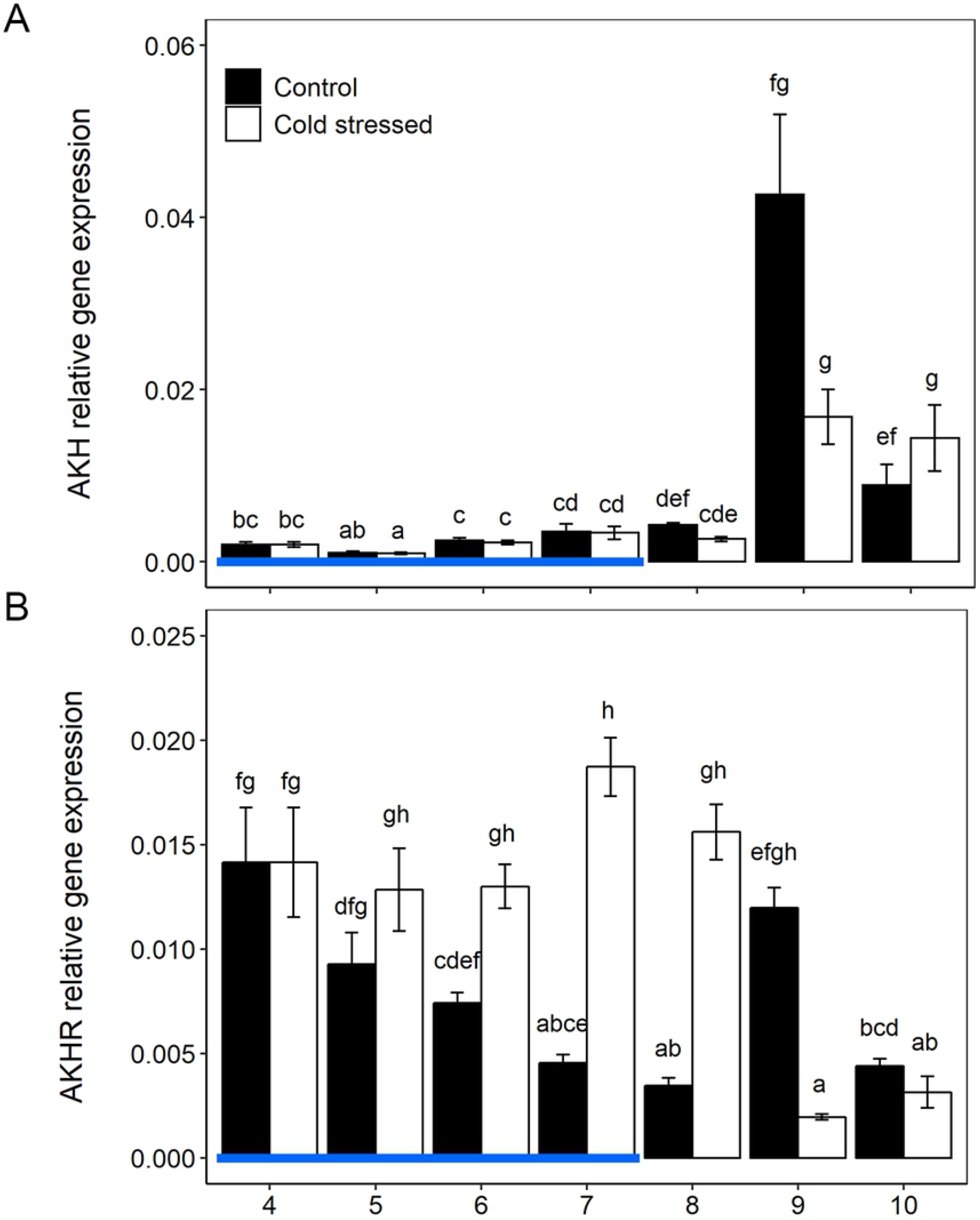
Transcript levels of components of the adipokinetic hormone (AKH) pathway in response to cold stress in honey bee brood. Four-day-old larvae were reared continuously at control conditions (34°C) or under cold stress (25°C) for 3 days and then were returned to control conditions. The relative gene expression of AKH (A) and AKH receptor (AKHR, B) was determined by RT-qPCR using Actin as a reference gene for normalization. The data are shown as the mean ± SD. Significant differences between control (black boxes) and stressed (white boxes) honey bee brood are indicated with different letters (ANOVA followed by post-hoc comparisons with Tukey). Blue line indicates the duration of cold stress.

S5 Fig shows that from day 4 to day 7 glucose and trehalose concentrations were not altered in the haemolymph of brood exposed to cold-stress compared to brood continuously reared at 34°C (Glucose: *t*_301_=−0.85, *P*=0.3944; Trehalose: *t*_140_=−0.24, *P*=0.8061; Fig S5).

## DISCUSSION

Here, we examined physiological and molecular responses in *A. mellifera* brood exposed to a short period of three days at low temperature. Firstly, we determined that growth rate is strongly diminished when larvae are incubated at 25°C (Fig 1A), and development progress is arrested (Fig 1B) compared to brood continuously reared at 34°C. When cold-stressed brood is returned to 34°C, the body mass is almost completely recovered in 24 h. Although delayed in time, we found that cold-stressed brood reaches the maximum value of body mass determined in brood reared at control temperature (S2B Fig), suggesting that compensatory mechanisms are operating. Compensatory growth is a response observed in organisms after a stressful period associated to decreased growth which allows recovering from the preceding delay once conditions improve (28). This can be achieved through augmented food consumption, increased metabolic efficiency, or altered developmental rate. In holometabolous insects, growth only occurs in the immature stages, so adult size is determined by the size the larva has reached when it stops feeding and begins metamorphosis (29). Size is an important indicator of fitness in insects (30). This reflects the relevance of understanding the molecular mechanisms and metabolic processes that determine the growth and development of honey bees during stressful conditions.

The first days after hatching from the egg, honey bee larvae have a high MR to sustain their rapid growth (31,32). Honey bee brood almost exclusively eat during larval stage to meet with the strong energy demand. Our results indicate that MR in honey bee brood reared at control temperature has a similar pattern to that reported by Schmolz and Lamprecht (31). However, we showed that during the period of cold-stress this pattern is severely affected (Fig 2). In insects under adverse environmental conditions, growth and development suppression is accompanied by a decrease in MR (33). We determined that 5- and 6-day-old cold-stressed brood display the same MR than brood growing at 34°C, indicating that probably a certain lag of time is required until MR is adjusted. Even more, in 7 and 8-days-old cold-stressed brood MR augmented compared to those reared at control temperature (Fig 2). In insects exposed to low temperature an increase in MR has been suggested to provide energy for cellular repairing processes (34). Thus, high levels of MR could be a necessary response contributing to the recovery process in cold-stressed honey bee brood.

We determined that ATP accumulates in honey bee brood reared continuously at 34°C from day 4 to day 8, and this correlated with a reduced MR in that period (S4 Fig). This could indicate that when brood reaches a final body size, growth ceases and energy must be stored for the metamorphosis (31). Furthermore, we showed that ATP content trend to diminish in cold-stressed brood transferred back to control temperature for one day. It would be interesting to further investigate this difference in ATP content to determine if cold-stressed brood consume their ATP for possible repair processes after stress.

On the other hand, the efficiency in the synthesis of ATP can vary significantly due to physiological conditions. Environmental temperature is one of the factors affecting the ATP production efficiency of organisms (35). UCPs decouple mitochondrial respiration from ATP synthesis, decreasing the oxidative phosphorylation efficiency (17) and promoting thermogenic processes (36). In invertebrates, however, scarce evidences have involved UCPs in heat generation. *D. melanogaster* larvae transferred from 23°C to 14°C for 1 h were 0.5°C warmer than the surrounding medium, suggesting that fly larvae could generate heat (25). By silencing *Ucp4C* the authors demonstrated that it is required for development through the prepupal stages at low temperatures. Furthermore, high activity of UCP in the fat body of *Gromphadorhina coquereliana* has been suggested to play a thermogenic role under cold-stress (37). Here, we showed that cold-stress strongly increase the transcript levels of MUP2 (Fig 3) and also we determined that ATP content trend to diminish in cold-stressed brood, together these results suggest a potential thermogenic role of MUP2 in honey bee brood reared at low temperatures. Moreover, in the hematophagous insect *Rhodnius prolixus*, upregulation of *UCP4* was accompanied by a decrease in reactive oxygen species (ROS) production suggesting that UCPs may act as a protective system for preventing oxidative stress damage after a blood meal (38). Thus, for better understanding of the possible role of MUP2 in cold-stressed honey bee brood further, detailed studies are required.

To get insights into the regulatory mechanisms that may be involved in honey bee brood exposed to cold-stress, two key metabolic pathways were evaluated: (i) insulin/insulin-like signalling (IIS) and (ii) adipokinetic hormone (AKH) (39). Transcript levels of *InR1* sharply increased in 5- and 6-days-old cold-stressed larvae exposed one or two days at 25°C, respectively (Fig 4D). High levels of InR1 would maximize the responsiveness to circulating ILP1. Previously, we demonstrated that *Insulin receptor substrate* (*IRS*), which functions in post-insulin receptor signal transduction, is highly expressed in 6-days-old cold-stressed brood. IIS pathway has been linked to increased resistance to cold stress (40), thus, elevated levels of their components may indicate a role in protection under cold stress in honey bee brood.

In well-fed *D. melanogaster* larvae, insulin producing cells of the brain secrete ILPs into the haemolymph and this is inhibited under poor nutrition. Once secreted, ILPs bind to the insulin receptor, activating IIS pathway and promoting cell growth. ILPs exert these effects, at least in part, through the FoxO transcription factor, a negative regulator of body growth (41). Here, we showed that *ILP2* is predominantly expressed in honey bee brood reared at 34°C compared to *ILP1*, as was reported (42). Additionally, we determined that *ILP2* transcript levels remain at low levels in cold-stressed brood until day 8 (Fig 4C). Interestingly, the high *ILP2* transcript levels coincide with the growth points at which the brood reached the maximum body mass, both in brood reared at control temperature and in brood exposed to cold-stress. When *D. melanogaster* reaches a critical weight this determines the duration of the growth period, and therefore the onset of metamorphosis (27). Therefore, ILP2 could be a key signal in honey bee brood mediating the developmental switch with the initiation of metamorphosis when optimal body mass is achieved. *FoxO* transcript levels remained low in honey bee brood independent of the rearing temperature but with a sharp peak in 7-days-old cold-stressed brood (Fig 4E). This result may indicate a role of FOXO arresting the development in cold-stressed brood and preventing them from continuing with the metamorphosis until optimal body mass and growth conditions are appropriate. Furthermore, in *Culex pipens*, shutdown of IIS prompts activation of FoxO, enhancing key characters essential for overwintering survival (43). Our results suggest that the IIS pathway coordinates developmental progress with brood growth, and particularly ILP2, InR1 and FoxO would play regulatory functions in response to cold-stress.

The AKH pathway can be involved in the regulation of carbohydrate metabolism in adult honey bee workers (44). Transcript levels of *AKH* were not altered during cold-stress in honey bee brood, while its receptor *AKHR* showed increased transcript accumulation all along the cold-stress period and even after returning to control temperature for one day (Fig 5). These results may indicate that the AKH pathway is turned on in honey bee brood during the period of cold exposure. We determined that glucose and trehalose levels in the haemolymph, which fuel MR, maintained stable in cold-stressed brood even though larvae reduced drastically food intake in that condition (S5 Fig). This result indicate that a mechanism would be activated to maintain glucose and trehalose homeostasis in cold stressed brood. In the antarctic midge *Belgica antarctica*, upregulation of genes supporting rapid glucose mobilization occurs during both heat and cold-stress (45). Our results support the hypothesis that the AKH pathway contributes to carbohydrate mobilization and homeostasis for sustaining basic energy requirements of an active metabolism in cold-stressed honey bee brood.

Overall, to compensate for the sharp drop in weight gain during adverse conditions, a fast growth is essential after the stress. Cold-stressed honey bee brood is able to increase one third of their body mass in one day after returning to regular growth temperature. An elevated MR in brood exposed to stress, as reported here, may provide energy for growth or cellular repairing processes as well as heat through mitochondrial decoupling mechanisms. In agreement with this, MUP2 transcripts were high in cold-stressed brood suggesting a role in thermogenic processes and/or in the attenuation of ROS formation. Fast expression changes in components of IIS and AKH pathways could regulate metabolic events for synchronizing growth and stress responses, and keep brood ready for a fast and strong response once normal nest temperature recovers. Our study improves the understanding of honey bee developmental biology under cold stress conditions being especially important in view of bee population decline.

## ACKNOWLEDGEMENTS

The authors are very grateful to Dr. Jesús D. Nuñez for his valuable help with the R software and Lic. Leonardo De Feudis for beekeeping support.

## SUPPORTING INFORMATION CAPTIONS

**S1 Fig.** Schematic diagram of the experimental design used to investigate physiological and molecular effects of cold stress in honeybee brood.

**S2 Fig.** Effect of cold stress on body mass gain during different phases of honeybee brood growth. (A) During the 3 days of exposure to 25°C. (B) In the maximum point of body mass acquisition in brood reared continuously at control temperature or under cold stress.

**S3 Fig.** Representative pictures of 7 and 8 days-old brood reared continuously at control (34°C) or exposed to cold stress at 25°C during 3 days. White brackets indicates the extension of food accumulated in the rearing well surrounding the larva.

**S4 Fig.** Changes in ATP content in honeybee brood under cold stress. Four-day-old larvae were reared continuously at control conditions (34°C) or under cold stress (25°C) for 3 days and then were returned to control conditions. ATP content was determined by a luminescent assay as was describe in Material and Methods. The data are shown as the mean ± standard deviation. Significant differences between honeybee brood incubated at control (black boxes) or cold stressed (white boxes) honeybee brood are indicated with different letters (ANOVA followed by post-hoc comparisons with Tukey). Blue line indicates the duration of cold stress.

**S5 Fig.** Glucose and trehalose concentrations in haemolymph of honeybee brood incubated under control or cold stress conditions.

